# A Computational Model of Phosphene Appearance for Epiretinal Prostheses

**DOI:** 10.1101/2021.06.07.447426

**Authors:** Jacob Granley, Michael Beyeler

## Abstract

Retinal neuroprostheses are the only FDA-approved treatment option for blinding degenerative diseases. A major outstanding challenge is to develop a computational model that can accurately predict the elicited visual percepts (phosphenes) across a wide range of electrical stimuli. Here we present a phenomenological model that predicts phosphene appearance as a function of stimulus amplitude, frequency, and pulse duration. The model uses a simulated map of nerve fiber bundles in the retina to produce phosphenes with accurate brightness, size, orientation, and elongation. We validate the model on psychophysical data from two independent studies, showing that it generalizes well to new data, even with different stimuli and on different electrodes. Whereas previous models focused on either spatial or temporal aspects of the elicited phosphenes in isolation, we describe a more comprehensive approach that is able to account for many reported visual effects. The model is designed to be flexible and extensible, and can be fit to data from a specific user. Overall this work is an important first step towards predicting visual outcomes in retinal prosthesis users across a wide range of stimuli.

## I. Introduction

Retinitis pigmentosa (RP) and age-related macular degeneration (AMD) are degenerative retinal diseases that lead to irreversible vision loss in more than 15 million people worldwide. As one promising treatment technology, retinal neuroprostheses [3], [19], [20], [24] aim to restore vision to these individuals by electrically stimulating the remaining retinal cells to evoke neural responses that are interpreted by the brain as visual percepts (“phosphenes”; see [2]).

However, a growing body of evidence suggests that the vision provided by these devices differs substantially from natural eyesight [7], [8], [10], [12]. Retinal implant users often report seeing distorted percepts and require extensive rehabilitative training to make use of their new vision [10]. Although single-electrode phosphenes are consistent from trial to trial, they vary across electrodes and users [7], [21]. Therefore, a deeper understanding of how electrical stimulation of the retina affects the quality of the generated artificial vision is crucial to designing more effective retinal implants.

A major outstanding challenge is to develop a computational model that can accurately predict visual outcomes for retinal implant users. Modeling the retinal response to electrical stimulation at a biophysical level (“bottom-up”) is challenging due to the complexity and variability of retinal circuitry in the presence of degeneration [18]. Even if an accurate user-specific biophysical model can be obtained, the detail required for simulation makes these methods too computationally expensive for many use cases [13].

In contrast, phosphene models are phenomenological (“top-down”) models constrained by behavioral data that predict visual perception directly from electrical stimuli [5]. To this end, Horsager *et al*. [16] predicted perceptual thresholds by convolving simulated pulse trains with a cascade of linear filters and nonlinear processing steps. Nanduri *et al*. [22] extended this model to generalize to suprathreshold stimulation. However, due to the number of free parameters and lack of an independent test set, these models should be viewed as descriptive rather than predictive models [22]. In addition, these models are unable to explain many reported spatial effects, such as phosphene elongation. To this end, Beyeler *et al*. [7] demonstrated that the phosphene shape elicited by epiretinal implants could be predicted by the spatial activation pattern of retinal nerve fiber bundles (NFBs). However, this cannot explain many reported temporal effects.

To address these challenges, we propose a phenomenological model constrained by both psychophysical [16], [22] and electrophysiological data [25] that predicts phosphene shape as a function of stimulus properties, such as amplitude, frequency, and pulse duration. The model is designed to be flexible and extensible so that it provides good predictions on average but can also be fit to data from a specific user.

## II. Methods

An overview of our model is given in Fig. 1. We assumed the subject is implanted with an epiretinal implant, such as Argus II [20] or POLYRETINA [11]. We focused on cathodic-first, square-wave, biphasic pulse trains, which make up the most common stimulus type in available devices. Given a stimulus, our model predicted the brightest “frame” of the percept seen by the user. Although the actual percept seen will likely grow and fade throughout the duration of stimulation, considering only the brightest frame made the problem tractable while allowing us to constrain the model with psychophysical data such as phosphene drawings and brightness ratings. A Python implementation based on pulse2percept [5] is available at https://github.com/bionicvisionlab/2021-BiphasicAxonMap.

**Fig. 1.**
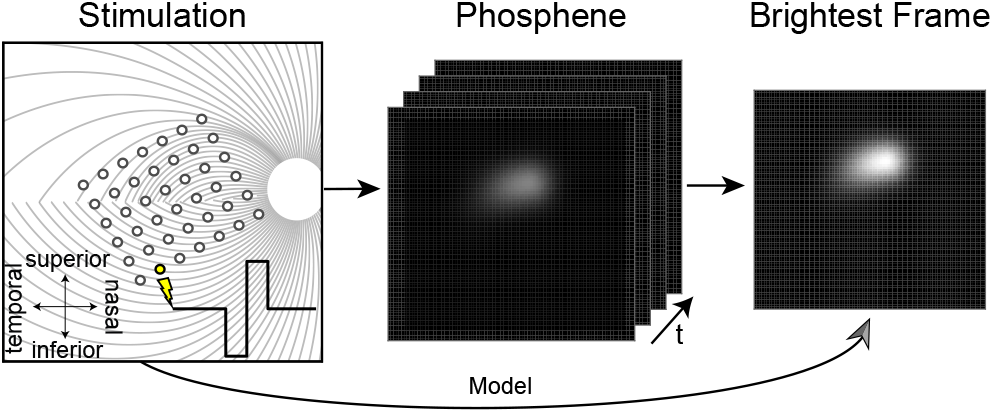
Phosphene model. A simulated biphasic pulse train is applied to a simulated epiretinal implant (*left*; for details see [5]), which evokes the percept of a flash of light whose brightness and shape evolve over time (*middle*). Rather than predicting the spatiotemporal properties of the elicited phosphene, the model directly predicts the brightest frame (*right*), which can be empirically validated using psychophysical data such as phosphene drawings and brightness ratings.

### A. Model Description

Our model extends the psychophysically validated axon map model [7] to account for a number of spatiotemporal effects. In this model, the shape of a phosphene generated by an epiretinal implant depended on the retinal location of the stimulating electrode. Because retinal ganglion cells (RGCs) send their axons on highly stereotyped pathways to the optic nerve [17], an electrode that stimulates nearby axons would antidromically activate RGC bodies located peripheral to the point of stimulation, leading to percepts that appear elongated in the direction of the underlying NFB trajectory (Fig. 2, *left*). The model assumed that an axon’s sensitivity to electrical stimulation decayed exponentially as a function of (i) distance from the stimulating electrode, with decay rate *ρ*, and (ii) distance along the axon from the cell body, with decay rate *λ* (Fig. 2, *right*). Since the maximum can occur either along the NFB or at the end of the NFB on the cell body, this model allows for both antidromic axonal and direct somatic RGC activation.

**Fig. 2.**
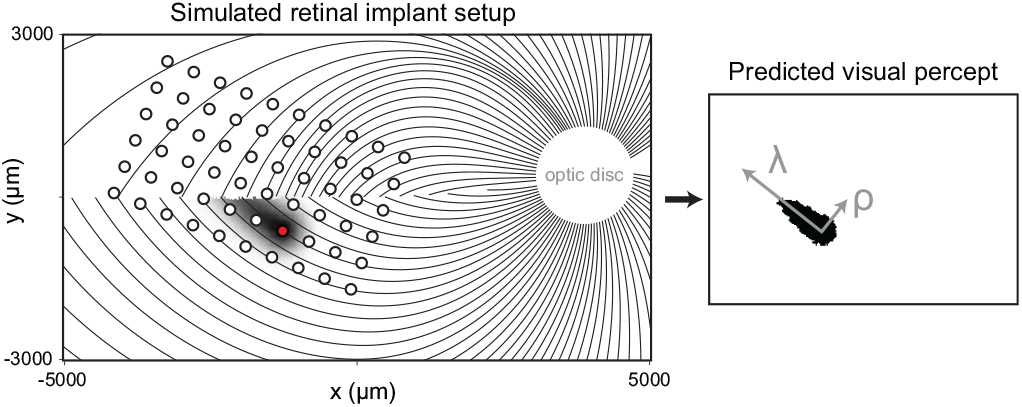
A simulated map of retinal NFBs (*left*) can account for visual percepts (*right*) elicited by retinal implants (reprinted with permission from [6]). *Left*: Electrical stimulation (red circle) of a NFB (black lines) could activate RGC bodies peripheral to the point of stimulation, leading to tissue activation (black shaded region) elongated along the NFB trajectory away from the optic disc (white circle). *Right*: The resulting visual percept appears elongated as well; its shape can be described by two parameters, *λ* (spatial extent along the NFB trajectory) and *ρ* (spatial extent perpendicular to the NFB). See Ref. [6] for more information.

We extended this model with three new terms (*F*_bright_, *F*_size_, and *F*_streak_; described in detail below) that controlled how a percept’s brightness, size, and streak length varied as a function of stimulus amplitude, frequency, and pulse duration. The output of the model was an intensity profile *I*(*r, θ*) that corresponded to the perceived brightness of a phosphene (polar coordinates centered over the fovea):

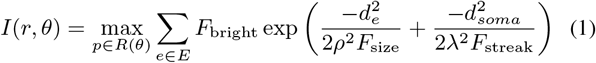

where *R*(*θ*) was the path of the axon to the point (*r, θ*), *p* was one individual point along the path, *d*_*e*_ was the Euclidean distance from *p* to the stimulating electrode, *E* was the set of all electrodes, and *d*_*soma*_ was the distance from *p* to the cell body along the axon, given by the path integral over the NFB:

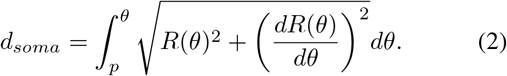

NFB paths (*R*(*θ*)) were modeled as spirals originating at the optic disc and terminating at each ganglion cell body [17]. The spirals were fit using manual axon fiber tracings of fundus images of 55 human eyes (for details see [17]).

Finding functions for *F*_bright_, *F*_size_, and *F*_streak_ that accurately describe phosphene appearance reported by retinal implant users is a crucial component of the model. Ideally, these functions would be fit to perceptual data from a specific user, where *F*_size_ would modulate phosphene size as a function of stimulus parameters, *F*_streak_ would modulate phosphene elongation, and *F*_bright_ would modulate overall phosphene brightness (see Eq. 1). However, obtaining the amount of data needed to fit such a model can be challenging, and any user-specific fit would be unlikely to generalize to other individuals. An alternative is therefore to fit a general model to data averaged across users.

### B. Model Fitting

Phosphene appearance is known to be affected by a number of stimulus parameters (Table I). Both Horsager *et al*. [16] and Nanduri *et al*. [22] found a linear relationship between stimulus frequency and brightness as well as amplitude and brightness for Argus I. Nanduri *et al*. also found a linear relationship between amplitude and size, and found no evidence for a relationship between frequency and size. Weitz *et al*. [25] analyzed the effect of varying pulse duration using *in vitro* mouse retina, and found inverse relationships between pulse duration and streak length, as well as pulse duration and threshold. To the best of our knowledge, no data has been published measuring the effect of amplitude and frequency on perceived phosphene streak length.

**TABLE I.**
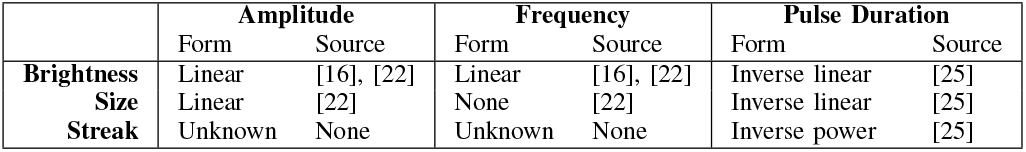
Stimulus parameters that affect phosphene appearance

To fit the data from these three different studies, we first converted raw amplitude values to a factor *a* of the threshold current for each individual electrode. Threshold was defined as the amplitude necessary to produce a visible percept with 50 % probability for a reference stimulus with the same frequency and a pulse duration of 0.45 ms. Since increases in pulse duration corresponded with increases in threshold amplitude [25], we needed to account for the resulting indirect effect on size and brightness by scaling amplitude *a* by a function of pulse duration, *t*:

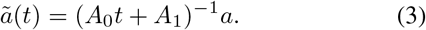

The final model was given by:

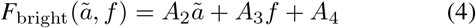

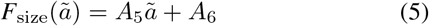

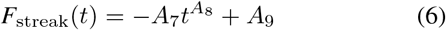

where *ã* was the scaled amplitude factor from Eq. 3, *f* was frequency in Hz, and *t* was pulse duration in ms. The scalars *A*_0_, …, *A*_9_ were the open parameters of the model (see code for values). All three equations were fit independently to data from the source papers using least squares regression.

Fig. 3 illustrates the model’s predicted percepts for single-electrode stimulation as a function of stimulation parameters. Percepts reflected a number of phenomena reported by epiretinal implant users. First, streaks were elongated along the underlying NFB [7]. Second, stimulus amplitude modulated phosphene size and brightness, but frequency only modulated brightness [22]. Third, if pulse duration were to increase with other parameters held constant, the brightness and size would decrease due to the threshold increasing [25], causing the percept to quickly dim and fade. Therefore, we also adjusted amplitude (Fig. 3, *bottom*) to offset the change in threshold, illustrating that longer pulse durations lead to shorter streaks.

**Fig. 3.**
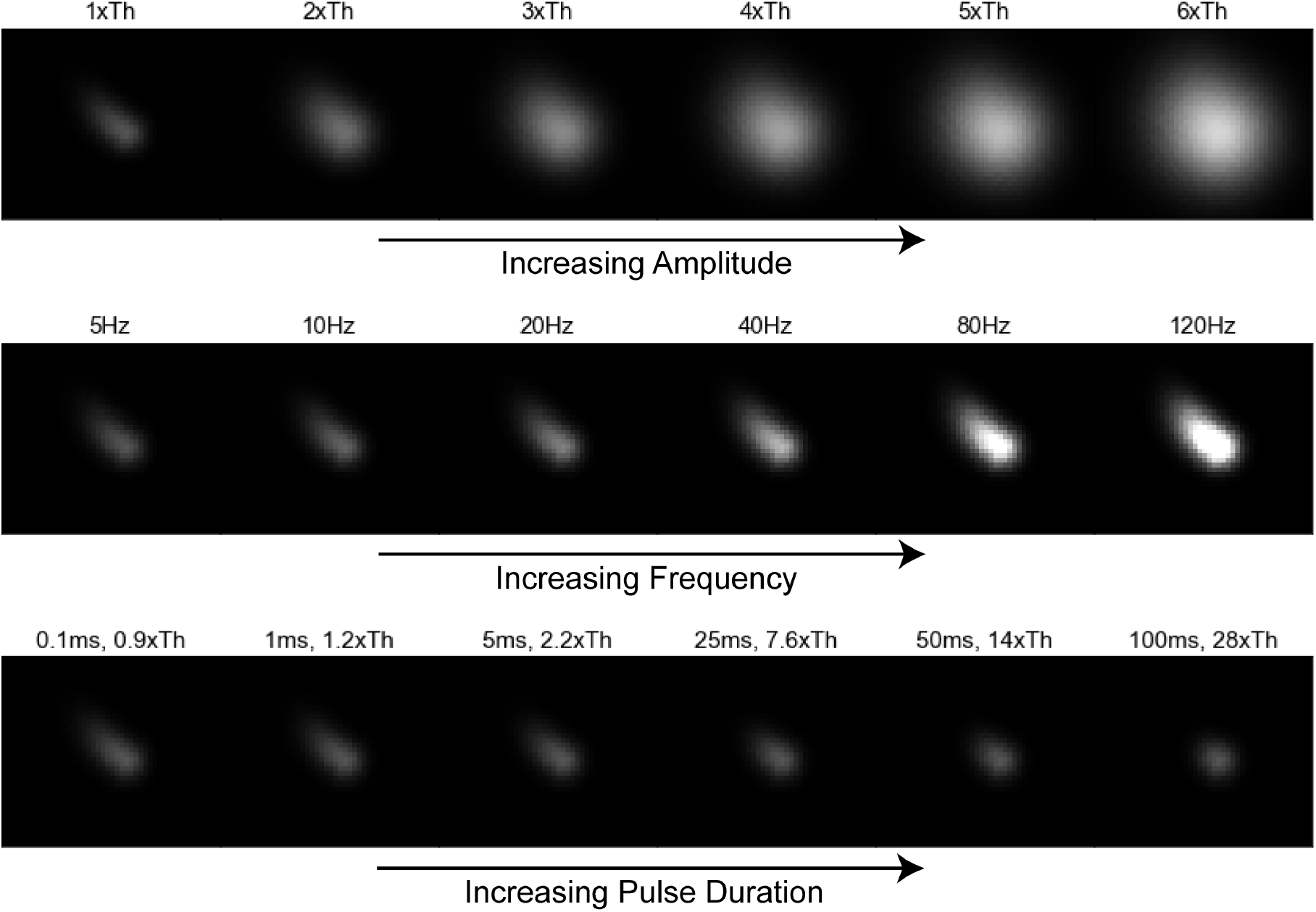
Predicted phosphene appearance as a function of amplitude (*top*), frequency (*middle*), and pulse duration (*bottom*). As pulse duration increased, we also increased the amplitude to offset the increase in threshold. The stimulating electrode was a disk electrode with radius 200 µm located in the central superior retina. Stimulus parameters not shown were kept constant (amplitude: 1xTh, frequency: 5 Hz, pulse duration: 0.45 ms, *λ*: 400 µm, *ρ*: 200 µm)

## III. Evaluation

In order to conduct a quantitative evaluation, we compared model predictions against data from two independent studies with epiretinal prosthesis users.

### A. Phosphene Appearance Across Stimulus Conditions

In the first experiment [22], an Argus I implant user was shown a reference stimulus as well as a test stimulus, which varied in either amplitude or frequency. The user was asked to rate the size or brightness of the test percept against the reference (Fig. 4, circles) on a scale from 0 to 100, with 10 meaning the brightness or size was equal to the reference, and 100 meaning 10x the reference.

**Fig. 4.**
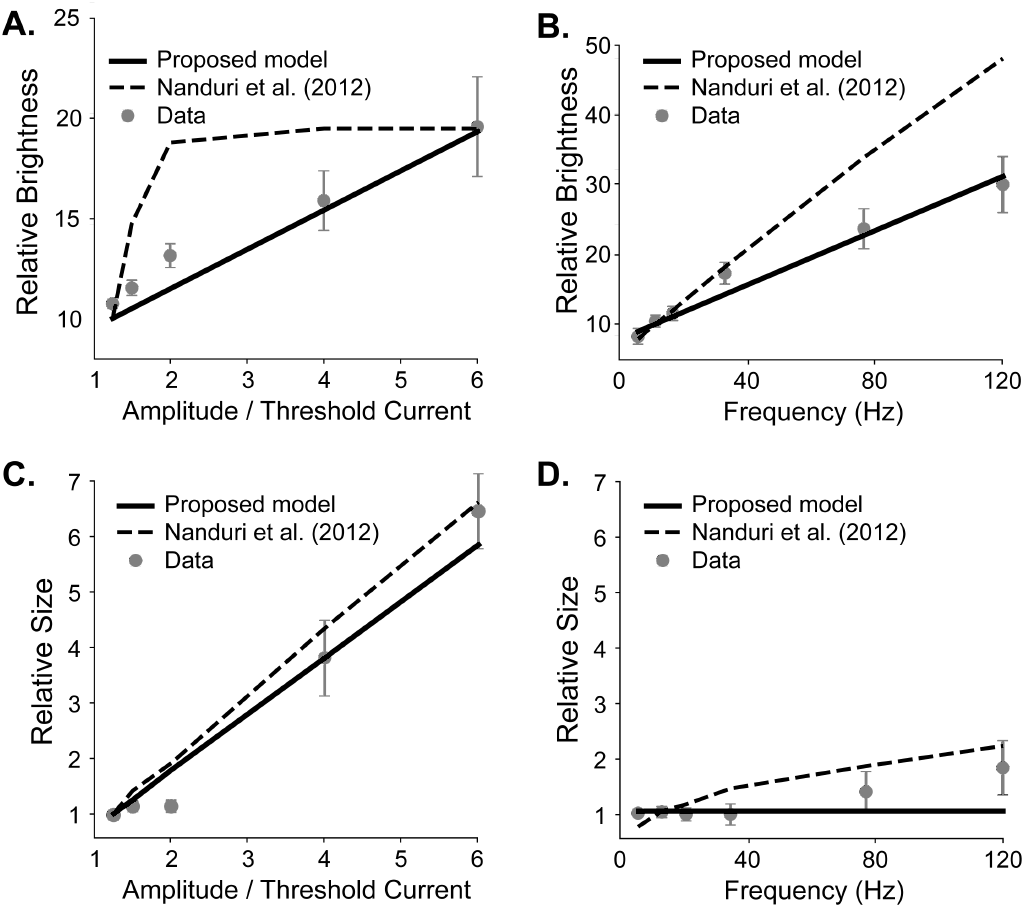
Brightness and size predictions for amplitude and frequency modulation on data from Nanduri *et al*. [22] (error bars: SEM). Solid lines: our model predictions. Dashed lines: Nanduri *et al*. model predictions.

To calculate predicted brightness, we ran both the reference and test stimuli through our model, and calculated the ratio between the predicted brightness of the test and reference pulse, multiplied by 10 to match the baseline in the psychophysical study. (Fig. 4A, B; solid black line). To estimate phosphene size, we counted the area with a predicted brightness greater than brightness at threshold, and again compared to the size of the reference stimulus (Fig. 4C, D; solid black line). Model predictions were compared to the baseline Nanduri model [22] (Fig. 4, dashed line).

Our model achieved a drastically better mean squared error (MSE) and *R*^2^ than the Nanduri model for predicting brightness with amplitude modulation (MSE: 0.9 vs 11.1, *R*^2^: 0.91 vs -0.07) and frequency modulation (MSE: 2.1 vs 71.9, *R*^2^: 0.97 vs -0.19), and marginally better measures for predicting size with amplitude modulation (MSE: 0.16 vs 0.2, *R*^2^: 0.965 vs 0.957). Additionally, our model predicted elongated phosphenes that matched user drawings, following the axon map model described in [7], whereas the Nanduri model always predicted circular percepts.

### B. Generalization to New Data

In order to evaluate the model’s ability to generalize, we tested on brightness rating data from Greenwald *et al*. [15] without refitting the model. The experiment conducted was a similar brightness rating task with two subjects, one being the same as in Nanduri *et al*. [22], but with different reference and test stimuli and electrodes.

The results are shown in Fig. 5. Here each data point is a single brightness rating on a particular electrode. The solid line shows our model’s predictions, the dashed line Nanduri *et al*. model predictions, while the dotted line depicts the linear model fit to the raw data using least squares. The Nanduri model performed poorly, showing it was unable to generalize beyond the data presented in [22]. Our model performed significantly better, obtaining an MSE that was slightly higher than the optimal linear model (model: 49.5, linear regression: 44.5, Nanduri *et al*.: 160.9), and a slightly worse *R*^2^ than the optimal linear model (model: 0.61, linear regression: 0.65, Nanduri *et al*.: -0.28).

**Fig. 5.**
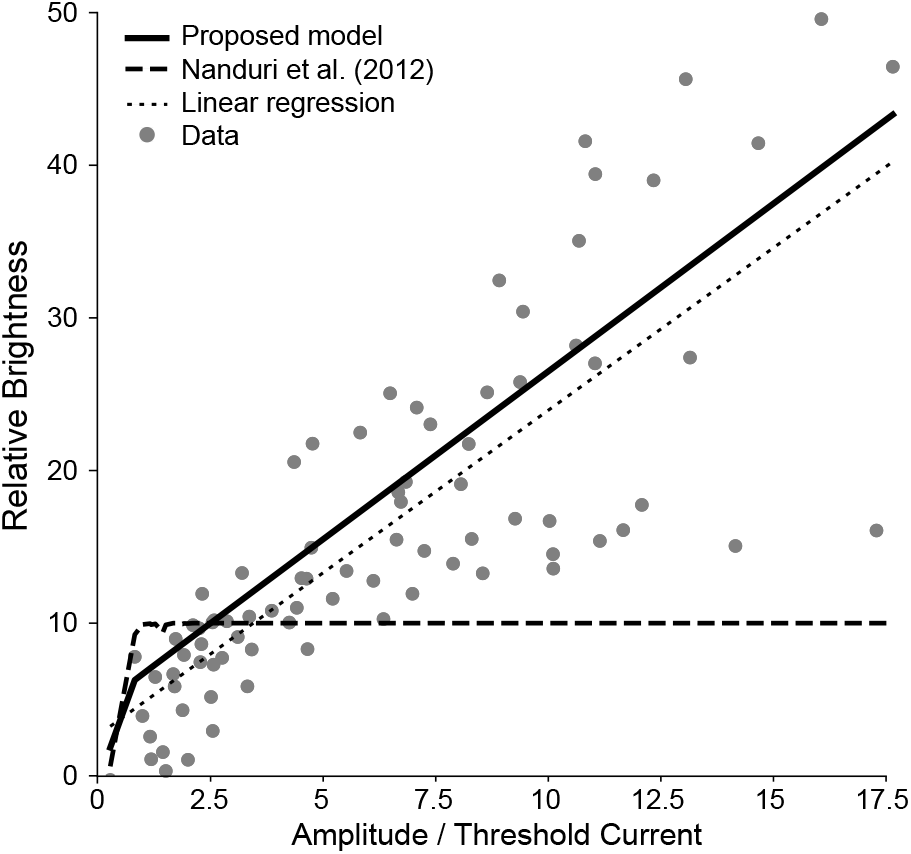
Brightness predictions for amplitude modulation on data from Greenwald *et al*. [15]. Solid line: our model prediction. Dashed line: Nanduri *et al*. predictions. Dotted line: Linear regression.

The stimuli in this experiment used different pulse durations, amplitude ranges, electrodes, and one different subject than the Nanduri *et al*. study, yet the model was able to give accurate predictions, even with its pulse duration equations being fit to neurophysiological data from mouse retina. These preliminary results suggest that our model may generalize well to new data, even for novel stimuli on new electrodes or implant users.

## IV. Discussion

We introduced a computational model constrained by both psychophysical and electrophysiological data that predicts phosphene appearance in epiretinal prostheses as a function of stimulus amplitude, frequency, and pulse duration. Whereas previous models focused on either spatial or temporal aspects of the elicited phosphenes, this is the first model able to account for all the listed effects. The model’s ability to accurately predict phosphene appearance for the new user introduced in the Greenwald *et al*. experiment [15] demonstrates the robustness of our model, and suggests it may work fairly well for new users as is. Overall this work is an important first step towards predicting visual outcomes in retinal prosthesis users across a wide range of stimuli.

Furthermore, our model is open-source, highly modular, and can easily be extended to account for additional effects if data is available. For example, if data is available on how other implant or stimulus properties affect phosphene appearance (*e*.*g*., stimulus waveform [14], electrode size [23], and electrode-retina distance [1]), Eqs. 3–6 could easily be updated to include the additional effects. As is, this work can be thought of as a general model that is able to make reasonable predictions on a patient-agnostic basis, similar in spirit to [16], [22]. However, since percepts vary widely across users, it is unlikely that any one model will be able to wholly describe every user’s unique perceptual experience. Future studies should thus focus on how this model could be fine-tuned to provide more accurate, patient-specific predictions.

In order to be used with a new user, Eqs. 3 - 6 should be refit to any brightness, size, and streak length ratings available from that user. Specifically, single-electrode phosphene drawings could be used to estimate patient-specific values for *ρ* and *λ* [7], and brightness ratings could be used to estimate *F*_bright_ [22]. However, little is known about how *F*_size_ (essentially a function of current spread in the tissue [9], [22]) and *F*_streak_ (a function of axonal activation [4], [25]) vary from patient to patient. With more data, future studies could streamline this process and identify the smallest set of experiments needed to fit a patient-specific model.

Although the present model is able to predict phosphene appearance across various stimulus conditions, there are a few limitations that should be addressed in future work. First, although Eqs. 3–6 are relatively simple, there are still ten free parameters in the model, which is comparable to the model described in [22]. However, the generalization performance of our model is noticeably improved. Second, lack of data prohibited a more thorough evaluation of our model. The dataset used for generalization contains brightness measurements for single-electrode stimuli from two subjects—but did not include data for phosphene size or streak length, nor for multi-electrode stimuli. Future work should therefore investigate the model’s ability to generalize across multi-electrode stimuli as well as different users and devices.

## V. Acknowledgments

This work was partially supported by the National Institutes of Health (NIH R00 EY-029329 to MB)

